# The palmitoyl acyltransferase DHHC5 mediates beta-adrenergic signaling in the heart by targeting Gα proteins and G protein-coupled receptor kinase 2

**DOI:** 10.1101/371211

**Authors:** Jessica J. Chen, Autumn N. Marsden, Askar M. Akimzhanov, Darren Boehning

## Abstract

S-palmitoylation is a reversible posttranslational modification that plays an important role in regulating protein localization, trafficking, and stability. Recent studies have shown that some proteins undergo extremely rapid palmitoylation/depalmitoylation cycles after cellular stimulation supporting a direct signaling role for this posttranslational modification. Here we investigated whether β-adrenergic stimulation of cardiomyocytes led to stimulus-dependent palmitoylation of downstream signaling proteins. We found that β-adrenergic stimulation led to increased Gαs and Gαi palmitoylation. The kinetics of palmitoylation was temporally consistent with the downstream production of cAMP and positive inotropic responses. Additionally, we identified for the first time that G protein-coupled receptor kinase 2 (GRK2) is a palmitoylated protein in cardiomyocytes. The kinetics of GRK2 palmitoylation were distinct from those observed with Gαs and Gαi after β-adrenergic stimulation. Knockdown of the plasma membrane-localized palmitoyl acyltransferase DHHC5 revealed that this enzyme is necessary for palmitoylation of Gαs, Gαi, and GRK2 and functional responses downstream of β-adrenergic stimulation. Our results reveal that DHHC5 activity is required for signaling downstream of β-adrenergic receptors.

## Introduction

Beta-adrenergic receptor (β-AR) signaling is one of the key regulators of contractile functions in the heart. Like other G protein-coupled receptors (GPCRs), β-ARs associate with the heterotrimeric G proteins, composed of α, β, and γ subunits (Gilman, 1987; Simon *et al.*, 1991). Upon activation, β-ARs act as a guanine nucleotide exchange factor and replace the GDP with GTP in the G protein α subunit (Hendriks-Balk *et al.*, 2008). Cardiac β-ARs associate with both Gαs and Gαi, which can stimulate or inhibit adenylyl cyclase, respectively (Xiao, 2001; Rockman *et al.*, 2002). Adenylyl cyclase produces the secondary messenger cAMP, which activates multiple downstream effectors, including protein kinase A (PKA) (Ishikawa *et al.*, 1992; Gottle *et al.*, 2009). PKA phosphorylates many substrates that alter contractile functions, including ryanodine receptors, the sarcoendoplasmic reticulum calcium ATPase (SERCA), and cardiac troponin I (Lindemann *et al.*, 1983; Takasago *et al.*, 1989; Valdivia *et al.*, 1997). Given the crucial roles of β-AR signaling in regulating cardiac function, it is not surprising that this pathway is tightly controlled. The β-adrenergic receptors are phosphorylated by GPCR kinases (GRKs) and are subsequently desensitized upon β-arrestin binding and downstream signaling (Inglese *et al.*, 1993; Barak *et al.*, 1999).

Multiple members of the β-AR signaling pathway undergo protein palmitoylation, a posttranslational modification that adds a 16-carbon palmitic acid to cysteine residues *via* a labile thioester bond. Protein palmitoylation is catalyzed by a class of enzymes termed palmitoyl acyltransferases (PATs). There are at least 23 mammalian PATs that share a conserved Asp-His-His-Cys (DHHC) domain that is essential for PAT activity (Fukata *et al.*, 2004). Most of the DHHC enzymes are localized to the Golgi apparatus and play crucial roles in the cycling of palmitoylated substrates between the Golgi apparatus and the plasma membrane. Recently our lab showed that the plasma membrane-localized DHHC21 is responsible for rapid, agonist-induced palmitoylation in T-cell signaling (Akimzhanov and Boehning, 2015).

Small G proteins such as Gα are classic palmitoylated proteins, and this modification is crucial for their plasma membrane targeting and subsequent functions (Linder *et al.*, 1993; Wedegaertner *et al.*, 1993). Studies in HeLa cells and primary hippocampal neurons suggested that Gα proteins can be palmitoylated by Golgi-localized DHHC3 and −7 (Tsutsumi *et al.*, 2009). The enzymatic activity regulating Gα protein palmitoylation in cardiac tissue is incompletely understood. Recently, the β_2_AR was reported to be palmitoylated at two cysteine residues in cardiomyocytes in a PKA-dependent manner resulting in internalization after sustained isoproterenol stimulation (Adachi *et al.*, 2016). In this study it was found that the Golgi-associated DHHC9, 14, and 18 may mediate this enzymatic activity. In order for palmitoylation to regulate GPCR signaling with kinetics consistent with a signaling role, presumably a plasma membrane localized DHHC enzyme should be activated. In cardiomyocytes the plasma membrane localized DHHC5 enzyme has been shown to be concentrated in caveolae and regulate the dynamic palmitoylation of phospholemman, a regulatory subunit of the Na pump (Tulloch *et al.*, 2011; Howie *et al.*, 2014). Importantly, phospholemman palmitoylation is dependent upon PKA phosphorylation, indicating it is likely palmitoylated in a stimulus-dependent manner (Tulloch *et al.*, 2011). Whether DHHC5 regulates the dynamic palmitoylation of other substrates after β-AR stimulation in cardiomyocytes is not known.

Here we show that Gαs, Gαi, and GRK2 are palmitoylated within minutes of β-AR activation in ventricular cardiomyocytes. The palmitoylation kinetics are consistent with the downstream cAMP production and changes in contractility. Knockdown of the plasma membrane-localized DHHC5 significantly inhibited agonist-induced palmitoylation and downstream responses indicating that this enzyme is required for β-AR signaling in cardiomyocytes.

## Results and Discussion

### Acyl-biotin exchange reveals palmitoylated proteins in the cardiac β-AR signaling pathway

Multiple methods have been developed to detect protein palmitoylation. The traditional method of metabolically labeling target proteins with [^3^H]-palmitate is powerful, yet is not ideal for unbiased identification of multiple palmitoylation substrates because the efficiency of labeling is dependent on the ability of cells to incorporate labeled palmitate and the palmitate turnover rate of each individual protein (Drisdel and Green, 2004). More recently, acyl-biotin exchange (ABE, Figure 1A) has been exploited as an efficient way of detecting the palmitoyl proteome in various tissues and organisms (Roth *et al.*, 2006; Kang *et al.*, 2008; Dowal *et al.*, 2011). The first step of ABE is blocking the free cysteine sites on proteins in cell lysates by using the thioreactive compound N-ethylmaleimide. Subsequently the labile thioester bond at the palmitoylation site is cleaved using neutral hydroxylamine exposing the free thiol group that can be subsequently biotinylated. After biotinylation, proteins are pulled down using streptavidin beads and eluted, followed by Western blotting to detect palmitoylated proteins. We used multiple rat cardiac cell types, including the myoblast cell line H9c2, neonatal rat ventricular myocytes (NRVM), and adult murine left ventricular tissue for detection of palmitoylated proteins downstream of β-AR (Figure 1B-D). We used calnexin as a positive control for protein palmitoylation in all experiments (Lynes *et al.*, 2012; Dallavilla *et al.*, 2016). As expected, we found that Gαs and Gαi are palmitoylated in both cell types and in cardiac tissue. Additionally, we also identified GRK2 as a novel palmitoylation target in H9c2 cells. The PKA regulatory subunit showed low levels of palmitoylation in H9c2 cells consistent with a previous proteomic study (Gould *et al.*, 2015). No palmitoylation was detected for other proteins downstream of β-ARs, including adenylyl cyclase 5/6 or PKA catalytic subunits (Figure 1B-D). Interestingly, although we could detect robust GRK2 palmitoylation in H9c2 cells (Figure 1B), we could not detect GRK2 palmitoylation in either NRVMs or cardiac tissue (Figure 1 C-D). We have previously shown Src kinases are palmitoylated in a stimulus-dependent manner (Akimzhanov and Boehning, 2015), and as such we hypothesized that GRK2 may be palmitoylated similarly in cardiomyocytes.

**Figure 1.**
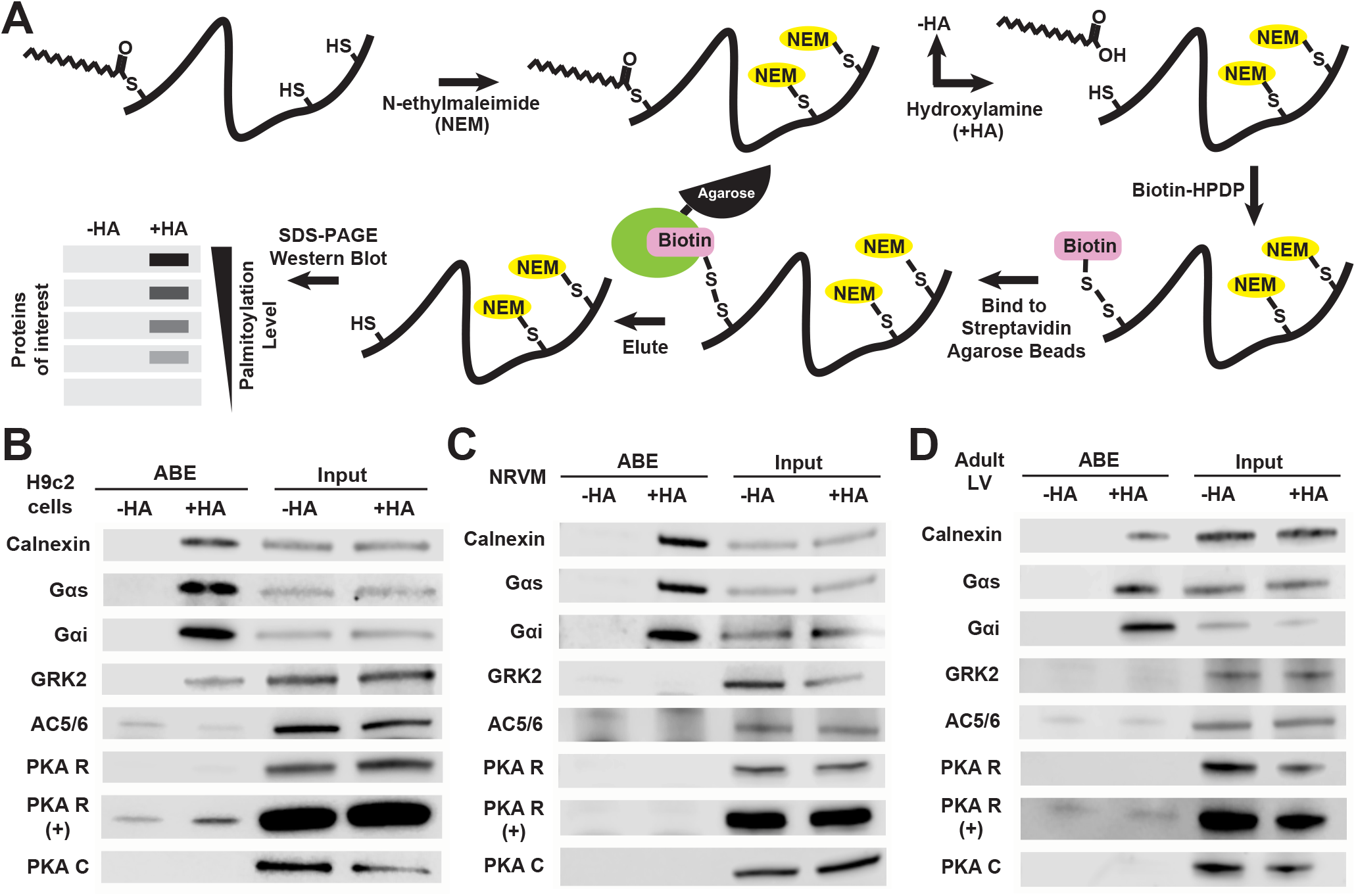
Identification of palmitoylated proteins in cardiomyocytes. (A) Schematic of the acyl biotin exchange (ABE) method used to quantify palmitoylation levels of cardiomyocytes proteins. (B) Palmitoylated proteins (ABE) and total lysate (Input) of indicated proteins in H9c2 cells. Reactions without hydroxylamine (-HA) are a negative control for non-specific binding to streptavidin agarose. Calnexin is used as a positive control for a palmitoylated protein. AC = adenylate cyclase; PKA R = PKA regulatory subunit; PKA C= PKA catalytic subunit. (+) indicates intentionally overexposed blot. (C-D) Same as in (B) except the ABE reaction was done on NRVM lysates (C) or murine left ventricular heart tissue (D).

### Activation of β-ARs induces rapid protein palmitoylation in cardiomyocytes

Previous studies in S49 mouse lymphoma cells and COS monkey kidney fibroblast-like cells suggested that stimulation of β-ARs increases the turnover of palmitate groups on Gα (Degtyarev *et al.*, 1993; Wedegaertner *et al.*, 1993). Additionally, though GRK2 has never been shown as a palmitoylated protein, another member of the GRK family GRK6 has been shown to be palmitoylated at the carboxyl-terminal domain in COS-7 cells (Stoffel *et al.*, 1994). To examine whether Gαs, Gαi, and GRK2 proteins are palmitoylated upon β-AR activation in cardiomyocytes we stimulated primary NRVMs with the β-AR agonist isoproterenol (ISO) and collected lysates from 0 to 30 minutes. We then performed ABE to quantify palmitoylation levels of proteins of interest as in Figure 1. We found that both Gαs and Gαi had significantly increased palmitoylation levels within a minute of ISO stimulation (Figure 2A,B). By 30 minutes the palmitoylation levels of Gαs and Gαi were significantly reduced suggesting a temporally distinct activation of acyl protein thioesterases. In contrast, the kinetics of GRK2 palmitoylation were characterized by increased levels at later time points compared to Gα proteins and were maintained throughout the 30 minute time course (Figure 2A-B). We next examined whether the agonist-induced palmitoylation is temporally consistent to downstream events mediating contractile functions. To test whether the kinetics of Gα palmitoylation correspond to downstream cAMP production, we measured intracellular cAMP levels in NRVMs after ISO stimulation. We found that cAMP levels increased within a minute of ISO stimulation and then decreased within 30min (Figure 2C), which was temporally consistent with the kinetics of Gα palmitoylation (Figure 2A,B). Isoproterenol is known to induce positive inotropy in NRVMs (Proven *et al.*, 2006). We used Fura-2 AM, a ratiometric calcium indicator, to monitor the intracellular calcium levels in spontaneously beating NRVMs. We observed an increase in calcium oscillation frequency (corresponding to beating frequency) within a minute of ISO stimulation, which like cAMP production, was consistent with the kinetics of Gα palmitoylation (Figure 2D). These results suggest that rapid palmitoylation of signaling proteins after β-AR stimulation may play a role in the assembly of the macromolecular β-AR signaling complex and efficient downstream modulation of cardiomyocyte contractility. To directly address this possibility, we targeted the enzymatic machinery potentially responsible for rapid palmitoylation of signaling proteins in cardiomyocytes.

**Figure 2.**
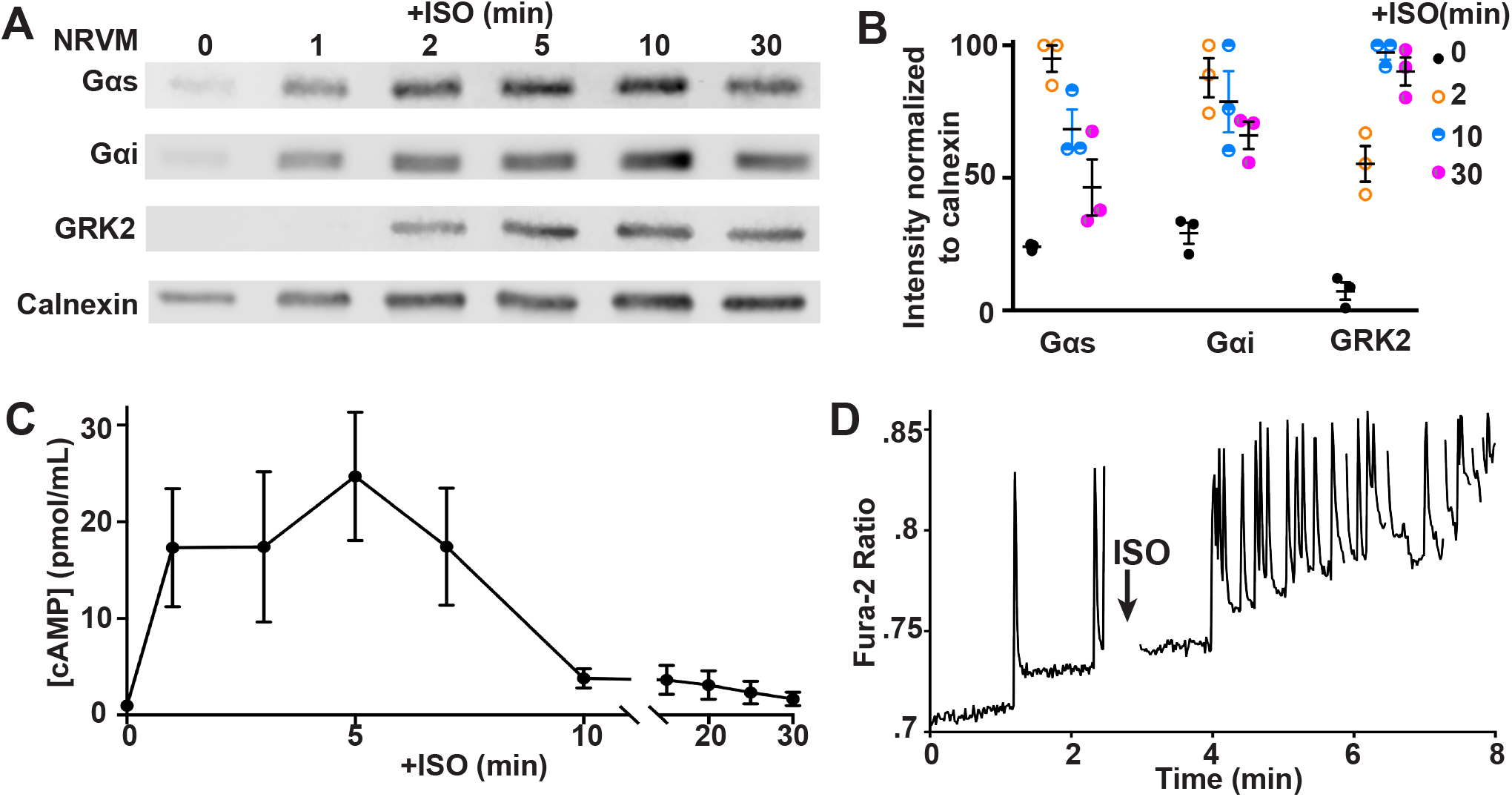
Isoproterenol induces rapid palmitoylation of Gαs, Gαi, and GRK2 in primary cardiomyocytes. (A) Neonatal rat ventricular myocytes (NRVMs) are treated with 10μM isoproterenol (ISO) for the indicated times. ABE was used to detect palmitoylation levels in response to ISO stimulation as in Figure 1. Calnexin, which has a relatively slow turnover of palmitoylation groups,was used to normalize turnover rates assessed by ABE. (B) Quantification of palmitoylation levels of Gαs, Gαi, and GRK2 normalized to calnexin at the indicated time points (n=3). (C) NRVMs were treated with 10μM ISO for indicated time points and intracellular cAMP concentration was measured using ELISA (n=5). (D) Representative trace of the Fura-2 ratio in a NRVM cell treated with 10μM ISO at the indicated time point.

### The palmitoyl acyltransferase DHHC5 mediates β-AR signaling in cardiomyocytes

We have previously shown that the plasma membrane-localized DHHC enzyme DHHC21 is responsible for rapid agonist-induced palmitoylation of signaling proteins in T cells (Akimzhanov and Boehning, 2015). Therefore, we hypothesized that the DHHC enzyme that catalyzes the ISO-induced palmitoylation in cardiomyocytes must similarly be localized to the plasma membrane. In addition to DHHC21, DHHC5 is one of the few DHHC enzymes that is plasma membrane localized (Ohno *et al.*, 2006). At the transcript level, DHHC5 is abundantly expressed in rat cardiomyocytes (Howie *et al.*, 2014). We analyzed the localization of DHHC5 in both the H9c2 rat myoblast cell line and NRVMs. Immunofluorescence staining showed that in both H9c2 and NRVMs, DHHC5 has a Golgi/ER localization (Figure 3). However, only in NRVMs is the plasma membrane localization of DHHC5 clearly evident, highlighting important differences between these two models (Figure 3B,C). To determine whether DHHC5 is responsible for ISO-induced palmitoylation in NRVMs, we used siRNA to knockdown DHHC5 in NRVMs (Figure 4A). We then performed ABE and found that agonist-induced palmitoylation of Gαs, Gαi, and GRK2 is reduced in DHHC5 knockdown cells (Figure 4B). Interestingly, basal levels of Gαs were increased suggesting a potential feedback loop between several DHHC enzymes to regulate Gαs palmitoylation (Figure 4B). Next we determined whether DHHC5 activity is required for signaling downstream of ISO stimulation in NRVMs. We found that DHHC5 knockdown cells have diminished levels of cAMP production following ISO stimulation (Figure 4C-D). Additionally, we found that DHHC5 knockdown cells no longer have an increased calcium oscillation frequency in response to ISO stimulation (Figure 4E-F). These data indicate that DHHC5 is essential for transmitting β-AR signaling in the heart, and this is likely mediated at least in part by regulating the palmitoylation state of Gαs, Gαi, and GRK2.

**Figure 3.**
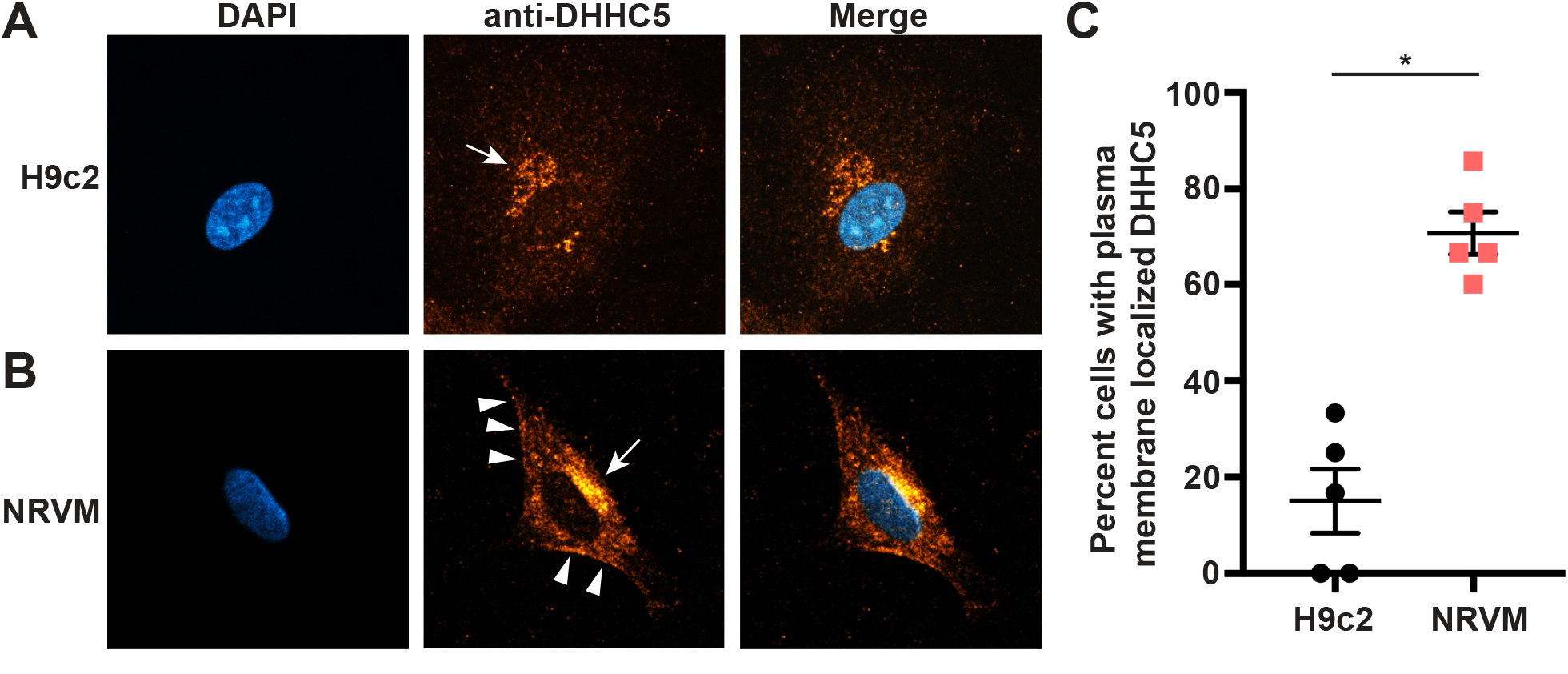
Localization of DHHC5 in H9c2 cells and primary cardiomyocytes. Immunofluorescence staining of H9c2 cells (A) and neonatal rat ventricular myocytes (B) with anti-DHHC5. Golgi apparatus localization is indicated by arrows, and plasma membrane localization is indicated by arrowheads. (C) Quantification of the percentage of cells with plasma membrane localized DHHC5 (n=5, * p=0.0001, unpaired T test).

**Figure 4.**
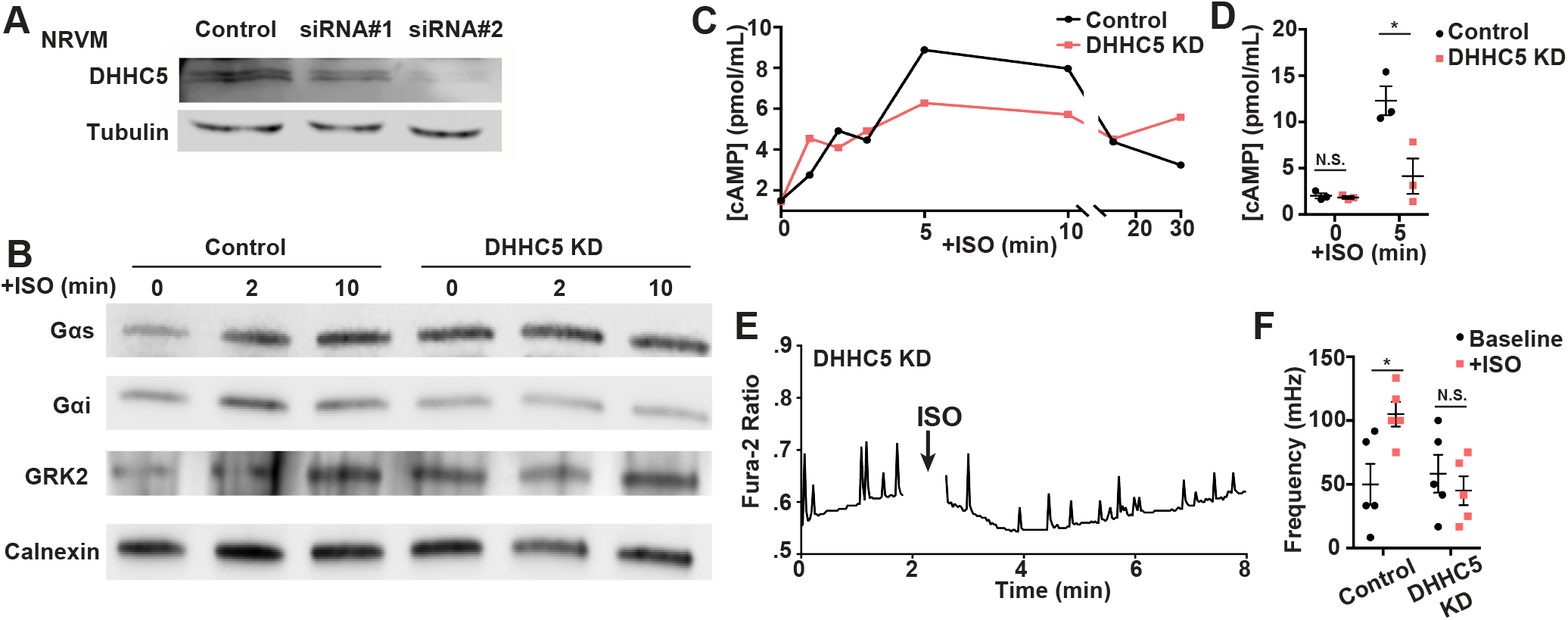
DHHC5 is required for signaling downstream of β-adrenergic receptor stimulation. (A) Western blot analysis of NRMVs transfected with control and two different siRNAs, both of which lead to reduced DHHC5 expression. Both siRNA are used in (B-F). (B) Palmitoylation of Gαs, Gαi, GRK2, and calnexin in control and DHHC5 knockdown NRVMs treated with 10μM isoproterenol (ISO) for indicated time points. (C) Cellular cAMP levels after ISO treatment in control (black) and DHHC5 knockdown (red) in NRVMs. (D) Quantification of cAMP levels before and after ISO (5min) averaged from three biological replicates. (*p=0.030; n.s. = not significant, p=0.60, paired T test). (E) Representative trace of the Fura-2 ratio in knockdown cells treated with ISO at indicated time point. (F) Quantification of beating frequency before (baseline) and after ISO (+ISO) in control and knockdown cells from five biological replicates (*p=0.018; n.s. = not significant, p=0.50, paired T test).

Our results show that DHHC5 is important for ISO-induced palmitoylation in cardiomyocytes. DHHC5 has a long C-terminal tail with a PDZ protein binding domain. One recent study suggested that the C-terminal tail of DHHC5 is important for its interactions with the substrate phospholemman in adult ventricular myocytes (Howie *et al.*, 2014). Additionally, in neurons phosphorylation of a tyrosine residue in the C-terminal tail of DHHC5 is key to its activity-regulated trafficking and subsequent substrate binding (Brigidi *et al.*, 2015). All of these studies suggested the importance of C-terminal tail in regulating DHHC5 activity. We propose that DHHC5 activity in the heart is likely to be tightly regulated, potentially by mechanisms related to β-AR signaling itself. One possibility is through PKA phosphorylation at Ser-704, part of a consensus PKA substrate motif KKXS, in the C-terminal tail. Therefore, future studies should be focused on elucidating the potential regulatory mechanisms that activate/inactivate DHHC5 in heart.

Very few studies have focused on the *in vivo* function of DHHC5 and its link to human diseases. One of the known substrates of DHHC5 is the neuronal scaffolding protein PSD-95 at the postsynaptic sites in neurons (Li *et al.*, 2010). Mice homozygotes for a hypomorphic allele of *zdhhc5* gene showed defects in learning and memory (Li *et al.*, 2010). Curiously, PDZ-95 palmitoylation levels are not decreased in the hypomorphic mice, likely because PDZ-95 can also be palmitoylated by other DHHC enzymes in neurons (Greaves *et al.*, 2011). The DHHC5 hypomorph mouse model has not been studied for potential cardiac defects, though half of the homozygotes are lost before birth, suggesting its role in early development (Li *et al.*, 2010). Most of the studies on cardiac functions of DHHC5 focus on its substrate phospholemman in regulating sodium pumps in adult ventricular myocytes (Tulloch *et al.*, 2011; Lin *et al.*, 2013; Howie *et al.*, 2014). Our data indicates that DHHC5 likely plays crucial role in β-AR signaling, a pathway that is closely associated with heart failure and target of multiple treatments.

In this study we also identified GRK2 as a novel palmitoylated protein in cardiomyocytes. A previous study showed that the GRK6 is palmitoylated at a cluster of three cysteine sites (Cys-562,562,565) in the carboxyl-terminal domain that is crucial for its plasma membrane localization (Stoffel *et al.*, 1994). However, this cluster of cysteine sites is not present in GRK2. Interestingly GRK2 is known to be isoprenylated, a non-reversible lipid modification, but isoprenylation alone is not sufficient for its plasma membrane targeting (Inglese *et al.*, 1992). Multiple studies have demonstrated that GRK2 directly binds the dissociated Gβγ subunits, which are also isoprenylated, and the formation of this complex is important for its plasma membrane localization and function (DebBurman *et al.*, 1996; Daaka *et al.*, 1997). Here we demonstrated that GRK2 proteins are only palmitoylated upon β-AR activation in cardiomyocytes, and this is potentially another way of efficiently targeting GRK2 to the plasma membrane. Future studies, including site-specific mutagenesis to identify the palmitoylation sites, are required to elucidate the functional importance of GRK2 palmitoylation in the heart.

## Materials and methods

### Antibodies, constructs, and reagents

Mouse monoclonal anti-Gαs was purchased from Millipore (MABN543). Rabbit monoclonal anti-Gαi/o and rabbit polyclonal anti-calnexin was purchased from Abcam (ab140125, ab22595). Rabbit polyclonal anti-GRK2, anti-PKA C-α, anti-PKA RI-α/β were purchased from Cell Signaling (#3982, 4782, 3927). Rabbit polyclonal anti-AC5/6 was purchased from Millipore (ABS573). Rabbit polyclonal anti-DHHC5 was purchased from Proteintech (21324-1-AP). Secondary antibodies conjugated to Alexa-488, Alexa-555 (Molecular Probes), peroxidase (Jackson ImmunoResaerch), IRDye 680LT, and 800CW (LI-COR) were used. Isoproterenol was purchased from Tocris (#1747) and was prepared fresh before use. Silencer Select DHHC5 siRNAs (s168613, s168614) and negative control medium GC duplex siRNA (12935112) were purchased from ThermoFisher Scientific. All other reagents were purchased at the highest possible purity from Sigma-Aldrich.

### Cell culture

The rat myoblast H9c2 cell line was obtained from American Type Culture Collection (ATCC) and maintained according to ATCC guidelines. Neonatal rat ventricular cardiomyocytes (NRVMs) were prepared from 1 to 2 day-old Sprague-Dawley rat pups, as previously described (Garcia *et al.*, 2017). Animals were purchased from Texas Animal Specialties (Humble, TX) and processed on the same day. Cardiomyocytes were plated on Primaria polystyrene plates (Corning) or fibronectin-coated glass coverslips. Cells were initially maintained in a mixture of 40% DMEM, 40% Ham’s F10 culture medium, and 20% FBS. Forty-eight hours after plating, the media was replaced by a mixture of 50% DMEM and 50% Ham’s F10 culture medium, supplied with human recombinant insulin (Gibco), transferrin, and thyroxine (Sigma). Cytosine-B-D-arabino-furanoside hydrochloride (10μM, Sigma) was present in all culture medium to prevent proliferation of fibroblasts or endothelial cells. Cardiomyocytes were transfected with Lipofectamine 3000 following manufacturer’s instruction. All experiments were carried out 48 hours following transfection and within one week of initial plating. The vertebrate animal procedures were approved by the Animal Welfare Committee (AWC) at UTHealth. For each assay, n represents number of biological replicates from different NRVM preps.

### Preparation of adult left ventricular lysates

Frozen left ventricle tissue from adult rats was a gift of Dr. Karmouty-Quintana (UTHealth) and Drs. Youker and Amione-Guerra (Houston Methodist Hospital). Frozen tissue was homogenized and lysed in 1% β-D-maltoside in PBS, supplemented with a protease inhibitor mixture (Roche 11873580001) and the acyl protein thioesterases inhibitor ML211 (10μM, Cayman Chemicals), followed by centrifugation (8000 ×g, 5min) to remove insoluble material. The cleared supernatant was stored at −80°C.

### Acyl biotin exchange

NRVMs were plated at a density of 1,000 cells per mm^2^ on 6-well plates. For control and DHHC5 knockdown experiments, the total amount of siRNA transfected was 30 pmol per 900,000 cells. NRVMs were stimulated with 10uM ISO at room temperature for indicated times, followed by scraping on ice and flash-freezing in liquid nitrogen. Cells were lysed in 1% β-D-maltoside in PBS, supplemented with a protease inhibitor mixture (Roche 11873580001) and acyl protein thioesterases inhibitor ML211 (10 μM, Cayman Chemicals). Acyl biotin exchange was performed as described previously with slight modification (Roth *et al.*, 2006). Briefly, proteins were precipitated by choloroform-methanol (CM) and incubated with 10mM N-ethylmaleimide (NEM) overnight at 4°C with gentle mixing. Following three rounds of CM precipitation, samples were incubated with 400 mM hydroxylamine (HA, pH=7, freshly prepared) and 1mM HPDP-biotin for 50 min at 37°C with gentle mixing. When a minus-HA treated sample was included, the sample was divided into two equal parts after the third precipitation and sodium chloride was used instead of HA. After three rounds of CM precipitation, samples were incubated with Streptavidin-agarose (Pierce) beads for at least 90 min up to 8 hrs at room temperature with gentle mixing. Following four rounds of washes, bound proteins were eluted for 15 min at 80°C with shaking in 30 μl of 1% β-mercaptoethanol and 2 mM DTT. The supernatant were transferred to new tubes and 20 μl of the sample were loaded onto a 4-20% gradient SDS-PAGE gel (BIO-RAD), followed by transfer to nitrocellulose membrane and immunoblotting.

### cAMP ELISA

For quantification of intracellular cAMP levels, NRVMs were plated at a density of 875 cells per mm^2^ on 24-well plates. Forty-eight hours following transfection, cells were stimulated with 1 μM ISO and assayed using the direct cAMP ELISA kit (Enzo) per manufacturer’s instructions. Each experiment was performed with three biological replicates.

### Calcium imaging

NRVMs were plated at a density of 300 cells per mm^2^ on fibronectin-coated glass coverslips. Cardiomyocytes were loaded with Fura-2 AM as described previously (Garcia *et al.*, 2017). Images were taken on a Nikon TiS inverted microscope with an image taken every second. Isoproterenol (10 μM) was added after 2 min of baseline recording. An oscillation was counted when the Fura-2 ratio rose 10% above the baseline ratio. Oscillation frequencies before and after ISO were calculated from 5 coverslips for each condition and mean ± SEM were plotted as described (Garcia *et al.*, 2017).

### Immunofluorescence staining

NRVMs were plated at a density of 250 cells per mm^2^ on fibronectin-coated glass coverslips. H9c2 cells were plated at the same density on uncoated glass coverslips. Immunofluorescence staining was performed as described previously (Garcia *et al.*, 2017). Anti-DHHC5 was used at a concentration of 1:100. For the quantification of plasma membrane localization of DHHC5, the percentage of cells displaying clear plasma membrane localization from five separate fields was averaged. The investigator was blinded to the cell type for these counts.

## Acknowledgements

We would like to thank Dr. Karmouty-Quintana (UTHealth) and Drs. Youker and Amione-Guerra (Houston Methodist Hospital) for kindly providing us with adult rat ventricular tissue. This work was supported by NIH grant R01GM081685 (DB) and AHA predoctoral fellowship 18PRE33960153 (JJC).

## Author contributions

J.J.C, A.A., and D.B. conceived the project. J.J.C. and A.M. performed the experiments. J.J.C., A.M., and D.B. analyzed data. All authors contributed to data interpretation and writing of the manuscript.

## Abbreviations

β-AR: Beta-adrenergic receptor
GRK: G protein-coupled receptor kinase
PKA: Protein Kinase A
PAT: palmitoyl acyltransferase
NRVM: neonatal rat ventricular myocytes
ABE: acyl-biotin exchange
HA: hydroxylamine
ISO: isoproterenol

## References

Adachi, N., Hess, D.T., McLaughlin, P., and Stamler, J.S. (2016). S-Palmitoylation of a Novel Site in the beta2-Adrenergic Receptor Associated with a Novel Intracellular Itinerary. J Biol Chem 291, 20232-20246.

Akimzhanov, A.M., and Boehning, D. (2015). Rapid and transient palmitoylation of the tyrosine kinase Lck mediates Fas signaling. Proc Natl Acad Sci U S A 112, 11876-11880.

Barak, L.S., Warabi, K., Feng, X., Caron, M.G., and Kwatra, M.M. (1999). Realtime visualization of the cellular redistribution of G protein-coupled receptor kinase 2 and beta-arrestin 2 during homologous desensitization of the substance P receptor. J Biol Chem 274, 7565-7569.

Brigidi, G.S., Santyr, B., Shimell, J., Jovellar, B., and Bamji, S.X. (2015). Activity-regulated trafficking of the palmitoyl-acyl transferase DHHC5. Nat Commun 6, 8200.

Daaka, Y., Pitcher, J.A., Richardson, M., Stoffel, R.H., Robishaw, J.D., and Lefkowitz, R.J. (1997). Receptor and G betagamma isoform-specific interactions with G protein-coupled receptor kinases. Proc Natl Acad Sci U S A 94, 2180-2185.

Dallavilla, T., Abrami, L., Sandoz, P.A., Savoglidis, G., Hatzimanikatis, V., and van der Goot, F.G. (2016). Model-Driven Understanding of Palmitoylation Dynamics: Regulated Acylation of the Endoplasmic Reticulum Chaperone Calnexin. PLoS computational biology 12, e1004774.

DebBurman, S.K., Ptasienski, J., Benovic, J.L., and Hosey, M.M. (1996). G protein-coupled receptor kinase GRK2 is a phospholipid-dependent enzyme that can be conditionally activated by G protein betagamma subunits. J Biol Chem 271, 22552-22562.

Degtyarev, M.Y., Spiegel, A.M., and Jones, T.L. (1993). The G protein alpha s subunit incorporates [3H]palmitic acid and mutation of cysteine-3 prevents this modification. Biochemistry 32, 8057-8061.

Dowal, L., Yang, W., Freeman, M.R., Steen, H., and Flaumenhaft, R. (2011). Proteomic analysis of palmitoylated platelet proteins. Blood 118, e62-73.

Drisdel, R.C., and Green, W.N. (2004). Labeling and quantifying sites of protein palmitoylation. Biotechniques 36, 276-285.

Fukata, M., Fukata, Y., Adesnik, H., Nicoll, R.A., and Bredt, D.S. (2004). Identification of PSD-95 palmitoylating enzymes. Neuron 44, 987-996.

Garcia, M.I., Karlstaedt, A., Chen, J.J., Amione-Guerra, J., Youker, K.A., Taegtmeyer, H., and Boehning, D. (2017). Functionally redundant control of cardiac hypertrophic signaling by inositol 1,4,5-trisphosphate receptors. J Mol Cell Cardiol 112, 95-103.

Gilman, A.G. (1987). G proteins: transducers of receptor-generated signals. Annu Rev Biochem 56, 615-649.

Gottle, M., Geduhn, J., Konig, B., Gille, A., Hocherl, K., and Seifert, R. (2009). Characterization of mouse heart adenylyl cyclase. J Pharmacol Exp Ther 329, 1156-1165.

Gould, N.S., Evans, P., Martinez-Acedo, P., Marino, S.M., Gladyshev, V.N., Carroll, K.S., and Ischiropoulos, H. (2015). Site-Specific Proteomic Mapping Identifies Selectively Modified Regulatory Cysteine Residues in Functionally Distinct Protein Networks. Chem Biol 22, 965-975.

Greaves, J., Carmichael, J.A., and Chamberlain, L.H. (2011). The palmitoyl transferase DHHC2 targets a dynamic membrane cycling pathway: regulation by a C-terminal domain. Mol Biol Cell 22, 1887-1895.

Hendriks-Balk, M.C., Peters, S.L., Michel, M.C., and Alewijnse, A.E. (2008). Regulation of G protein-coupled receptor signalling: focus on the cardiovascular system and regulator of G protein signalling proteins. Eur J Pharmacol 585, 278-291.

Howie, J., Reilly, L., Fraser, N.J., Vlachaki Walker, J.M., Wypijewski, K.J., Ashford, M.L., Calaghan, S.C., McClafferty, H., Tian, L., Shipston, M.J., Boguslavskyi, A., Shattock, M.J., and Fuller, W. (2014). Substrate recognition by the cell surface palmitoyl transferase DHHC5. Proc Natl Acad Sci U S A 111, 17534-17539.

Inglese, J., Freedman, N.J., Koch, W.J., and Lefkowitz, R.J. (1993). Structure and mechanism of the G protein-coupled receptor kinases. J Biol Chem 268, 23735-23738.

Inglese, J., Koch, W.J., Caron, M.G., and Lefkowitz, R.J. (1992). Isoprenylation in regulation of signal transduction by G-protein-coupled receptor kinases. Nature 359, 147-150.

Ishikawa, Y., Katsushika, S., Chen, L., Halnon, N.J., Kawabe, J., and Homcy, C.J. (1992). Isolation and characterization of a novel cardiac adenylylcyclase cDNA. J Biol Chem 267, 13553-13557.

Kang, R., Wan, J., Arstikaitis, P., Takahashi, H., Huang, K., Bailey, A.O., Thompson, J.X., Roth, A.F., Drisdel, R.C., Mastro, R., Green, W.N., Yates, J.R., 3rd, Davis, N.G., and El-Husseini, A. (2008). Neural palmitoyl-proteomics reveals dynamic synaptic palmitoylation. Nature 456, 904-909.

Li, Y., Hu, J., Hofer, K., Wong, A.M., Cooper, J.D., Birnbaum, S.G., Hammer, R.E., and Hofmann, S.L. (2010). DHHC5 interacts with PDZ domain 3 of postsynaptic density-95 (PSD-95) protein and plays a role in learning and memory. J Biol Chem 285, 13022-13031.

Lin, M.J., Fine, M., Lu, J.Y., Hofmann, S.L., Frazier, G., and Hilgemann, D.W. (2013). Massive palmitoylation-dependent endocytosis during reoxygenation of anoxic cardiac muscle. Elife 2, e01295.

Lindemann, J.P., Jones, L.R., Hathaway, D.R., Henry, B.G., and Watanabe, A.M. (1983). beta-Adrenergic stimulation of phospholamban phosphorylation and Ca2+-ATPase activity in guinea pig ventricles. J Biol Chem 258, 464-471.

Linder, M.E., Middleton, P., Hepler, J.R., Taussig, R., Gilman, A.G., and Mumby, S.M. (1993). Lipid modifications of G proteins: alpha subunits are palmitoylated. Proc Natl Acad Sci U S A 90, 3675-3679.

Lynes, E.M., Bui, M., Yap, M.C., Benson, M.D., Schneider, B., Ellgaard, L., Berthiaume, L.G., and Simmen, T. (2012). Palmitoylated TMX and calnexin target to the mitochondria-associated membrane. EMBO J 31, 457-470.

Ohno, Y., Kihara, A., Sano, T., and Igarashi, Y. (2006). Intracellular localization and tissue-specific distribution of human and yeast DHHC cysteine-rich domain-containing proteins. Biochim Biophys Acta 1761, 474-483.

Proven, A., Roderick, H.L., Conway, S.J., Berridge, M.J., Horton, J.K., Capper, S.J., and Bootman, M.D. (2006). Inositol 1,4,5-trisphosphate supports the arrhythmogenic action of endothelin-1 on ventricular cardiac myocytes. J Cell Sci 119, 3363-3375.

Rockman, H.A., Koch, W.J., and Lefkowitz, R.J. (2002). Seven-transmembrane-spanning receptors and heart function. Nature 415, 206-212.

Roth, A.F., Wan, J., Bailey, A.O., Sun, B., Kuchar, J.A., Green, W.N., Phinney, B.S., Yates, J.R., 3rd, and Davis, N.G. (2006). Global analysis of protein palmitoylation in yeast. Cell 125, 1003-1013.

Simon, M.I., Strathmann, M.P., and Gautam, N. (1991). Diversity of G proteins in signal transduction. Science 252, 802-808.

Stoffel, R.H., Randall, R.R., Premont, R.T., Lefkowitz, R.J., and Inglese, J. (1994). Palmitoylation of G protein-coupled receptor kinase, GRK6. Lipid modification diversity in the GRK family. J Biol Chem 269, 27791-27794.

Takasago, T., Imagawa, T., and Shigekawa, M. (1989). Phosphorylation of the cardiac ryanodine receptor by cAMP-dependent protein kinase. J Biochem 106, 872-877.

Tsutsumi, R., Fukata, Y., Noritake, J., Iwanaga, T., Perez, F., and Fukata, M. (2009). Identification of G protein alpha subunit-palmitoylating enzyme. Mol Cell Biol 29, 435-447.

Tulloch, L.B., Howie, J., Wypijewski, K.J., Wilson, C.R., Bernard, W.G., Shattock, M.J., and Fuller, W. (2011). The inhibitory effect of phospholemman on the sodium pump requires its palmitoylation. J Biol Chem 286, 36020-36031.

Valdivia, L.A., Sun, H., Rao, A.S., Tsugita, M., Chen, C.T., Park, I.Y., Fung, J.J., and Starzl, T.E. (1997). Donor-specific transfusion in the nude rat prolongs survival of subsequently transplanted hamster cardiac xenografts. Transplant Proc 29, 928-929.

Wedegaertner, P.B., Chu, D.H., Wilson, P.T., Levis, M.J., and Bourne, H.R. (1993). Palmitoylation is required for signaling functions and membrane attachment of Gq alpha and Gs alpha. J Biol Chem 268, 25001-25008.

Xiao, R.P. (2001). Beta-adrenergic signaling in the heart: dual coupling of the beta2-adrenergic receptor to G(s) and G(i) proteins. Sci STKE 2001, re15.

